# The Effect of Attention Load on Balance Control Performance

**DOI:** 10.1101/2021.03.28.437397

**Authors:** Sheida Mirloo, Behzad Parpanchi, Golnaz Baghdadi

## Abstract

Posture balance control is an essential ability that is affected by the attention load. We investigated the effect of attention load on posture balance control experimentally and computationally. Fifteen young individuals participated in an experiment containing simultaneous performing of a balance control task and an auditory task. A previous computational model was extended by introducing the effect of attention load as a gain in a Proportional-Integral-Derivative (PID) controller. Results demonstrated that the sensitivity of the posture balance control to the attention load should be considered besides other influential factors in designing sport or physical rehabilitation exercises. Simulations suggested that the issues of joint impedance stiffness or viscosity might also be compensated by changing the attention load.

## 1. Introduction

Control of the body position in space to attain desirable balance and orientation is considered as the definition of posture balance control (Woollacott & Shumway-Cook, 2002). Posture balance control is also defined as the ability of one to maintain the center of pressure (CoP) within the stability boundary (Trapp, 2013). The CoP is referred to as the point that the earth’s vertical reaction force vector is applied (Winter, Patla, & Frank, 1990). The proper positions of both CoP and the center of mass (CoM) have a vital role in balance control. CoM is referred to as a point which is equivalent to the total mass of the body in the global reference system (GRS) and is the weighted mean of the mass center of each body section in three-dimensional space (Winter, 1995). The distance between CoP and CoM is suggested as a variable that is sensitive to perturbations and stability while standing (van der Kooij, van Asseldonk, & van der Helm, 2005). The posture balance control system can be divided into the central nervous system (CNS) and the muscular-skeleton system (Winter et al., 1990).

The importance of balance control has encouraged many researchers to investigate the different factors that can enhance or disrupt it. Although balance control was traditionally considered as an automatic task, recently, it was found out that balance control demands attention (Trapp, 2013).

Attention is explained as the individuals’ capacity to process information, and it can be considered as the capability of focus and selective environment information processing while ignoring distractors (McCulloch, Buxton, Hackney, & Lowers, 2010b). One of the accepted theories of attention is that the capacity of information processing for each individual is limited, and each task needs some of this capacity. Therefore, if two tasks are done simultaneously, and they both need a high amount of attention, the performance in both or at least one of the tasks will reduce (Woollacott & Shumway-Cook, 2002). The goal of the attention is to bias the competition towards a goal or an expected event (Chun, Golomb, & Turk-Browne, 2011). It was shown that simultaneous performing of cognitive and balance control tasks could disrupt the performance in both tasks (Mahboobin, Loughlin, & Redfern, 2007; Woollacott & Shumway-Cook, 2002) (for more details, see references listed in Table A.1 in Appendix). Balance maintenance impairment due to the improper attention resource allocation in elderly people is another example that shows the relationship between attention and balance control (Brauer, Woollacott, & Shumway-Cook, 2002). Most of the studies have shown that the division of attention leads to an increase in body fluctuations around its axis of balance (Hyndman & Ashburn, 2003). The level of attention demanded by a balance control task depends on age, balance capability, accessibility to sensory information, the difficulty of the balance task, impairments of the body, and the power of processing of information in individuals (Geurts & Mulder, 1994; Hyndman, Ashburn, Yardley, & Stack, 2006; Shorer, Becker, Jacobi-Polishook, Oddsson, & Melzer, 2012; Woollacott & Shumway-Cook, 2002). In previous studies, the relationship between attention and balance control was investigated experimentally. Since this relationship has not fully known yet, we decided to investigate it further using the power of computational models. We also performed a new experiment to overcome some drawbacks of previous ones. In previous studies, the cognitive tasks, which were performed by individuals during the balance control task, demanded individuals’ perception speed, calculation capability, or some other cognitive resources that could affect the final performance due to their variability between individuals. To avoid this issue, we tried to investigate the role of attention using a straightforward dual-task experiment, which did not activate complicated cognitive functions. Therefore, we could reduce individual variability in doing attention demanded tasks. The experiment includes a balance control task that was simultaneously performed with an auditory-oral response task.

Another novelty of the current study is introducing a new output criterion to investigate the effect of attention on balance control moreover than the reaction time, which is usually used in similar studies. The new criterion is based on the fluctuations of the board, which is placed under the feet of the participants in the balance control task.

Various models have been proposed for postural or dynamic balance control (Kuo, 1995, 2005; Masani, Vette, & Popovic, 2006) without considering the role of attention. In the current study, a new computational model was also proposed by modifying the previously suggested model to add the role of attention.

## 2. Materials and Methods

### 2.1. Subjects

Fifteen healthy right-handed female volunteers in the age range of 20 – 25 (median=22) participated in our study. They had no history of neurological or motor disorders. The experiment was performed under the approval of the Biomedical Engineering Department of the Amirkabir University of Technology, Tehran, Iran (#10115/A/173). All the participants were aware of the required parts of the experiment procedure and signed the informed consent form. Before the beginning of the experiment, participants filled a questionnaire to score their concentration, mental fatigue, and amounts of sleep in the last 24 hours. We used the results of this questionnaire to exclude participants who were so tired or did not have enough concentration. We also exclude individuals who were professional or had some experience in some sports that need posture balance, such as gymnastics. These exclusions were because fatigue and lack of concentration or an extraordinary ability to maintain balance can affect the interpretation of results.

### 2.2. Experiment

The experiment had two stages. In the first stage, participants were standing on a wobble board while trying to keep their posture balance and stability (i.e., single task). In the second stage, participants were trying to keep their posture balance and stability on the wobble board while they were listening to audio numbers that were played in their left/right ear (i.e., dual-task). After the presentation of each audio number, individuals were asked to say what number they heard from which ear. For instance, if the number “16” is played in the “left” ear, the individual will be expected to say, “Sixteen left.” Therefore, in the first stage, individuals’ focus was on keeping the posture balance, while in the second stage, their focus was divided between balance controlling and processing auditory stimuli. Due to the division of attention, less attention was attributed to the posture balance controlling. Hence, comparing the posture balance performance of individuals in two stages can reveal the role of attention in the posture-balance-control system. It should be noted that before the experiment, participants performed a short training stage to be familiar with the experiment setup.

The second stage consisted of 61 trials in each of which a number was played pseudorandomly into the left or the right ear. Auditory stimuli were played for 500 milliseconds with an intensity of 100 db through a headphone. The inter-stimulus interval was 2.5 seconds.

To quantify the performance of participants in keeping their posture balance, we measured the amount of wobble board deviation from a horizontal line shown in Figure 1.b (the red line). Measuring the deviation was performed by inspecting images that were taken from the wobble board during the experiment through a webcam, which was fixed in front of the wobble board (Figure 1.a). Regardless of the role of attention, talking and naming the numbers themselves can cause imbalance. Therefore, measurement of deviation was performed in the time of quiet standing (while participants were only listening to the numbers).

**Figure 1.**
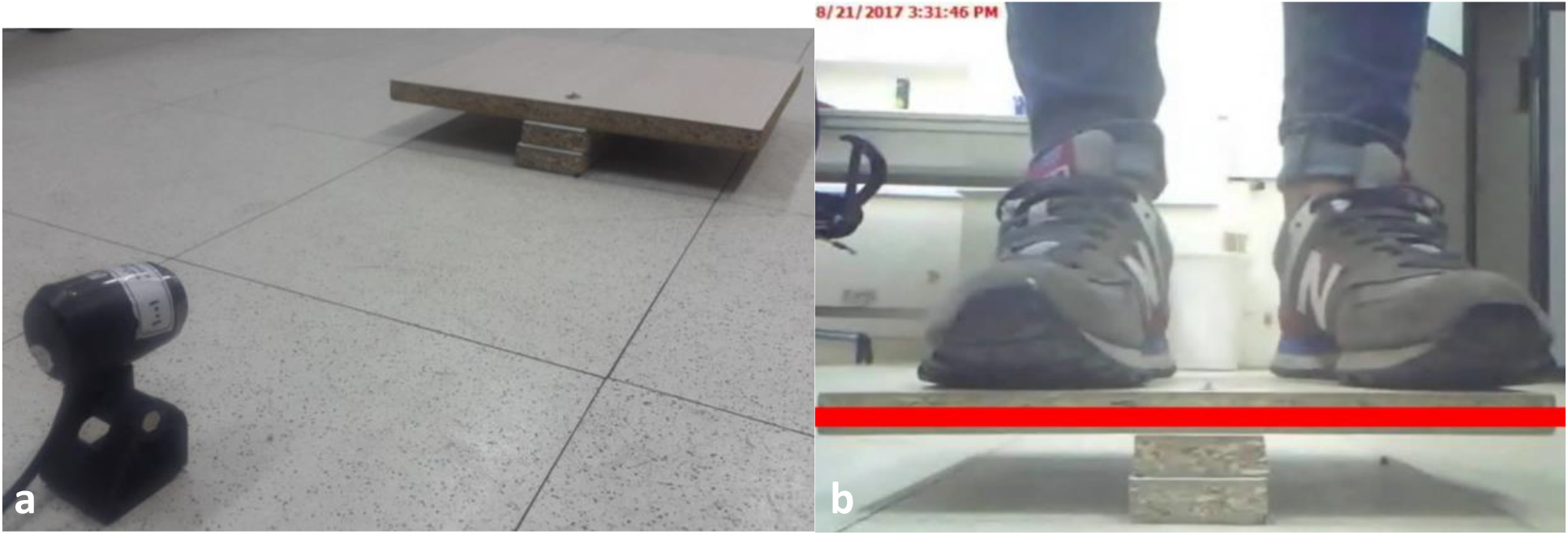
a. Experiment setup: a webcam placed in front of the wobble board. b. A representation of an individual standing on the wobble board while she is keeping her posture balance (deviation of the wobble board from a horizontal red line demonstrates the posture balance control performance)

### 2.3. The suggested model

In this study, we extended the model suggested by Masani et al. (2006) (Masani et al., 2006). We added a block to this model to simulate the effect of attention. As shown in Figure 2, an inverted pendulum model represents the body’s dynamic and kinematic (Eq. (1)).

**Figure 2.**
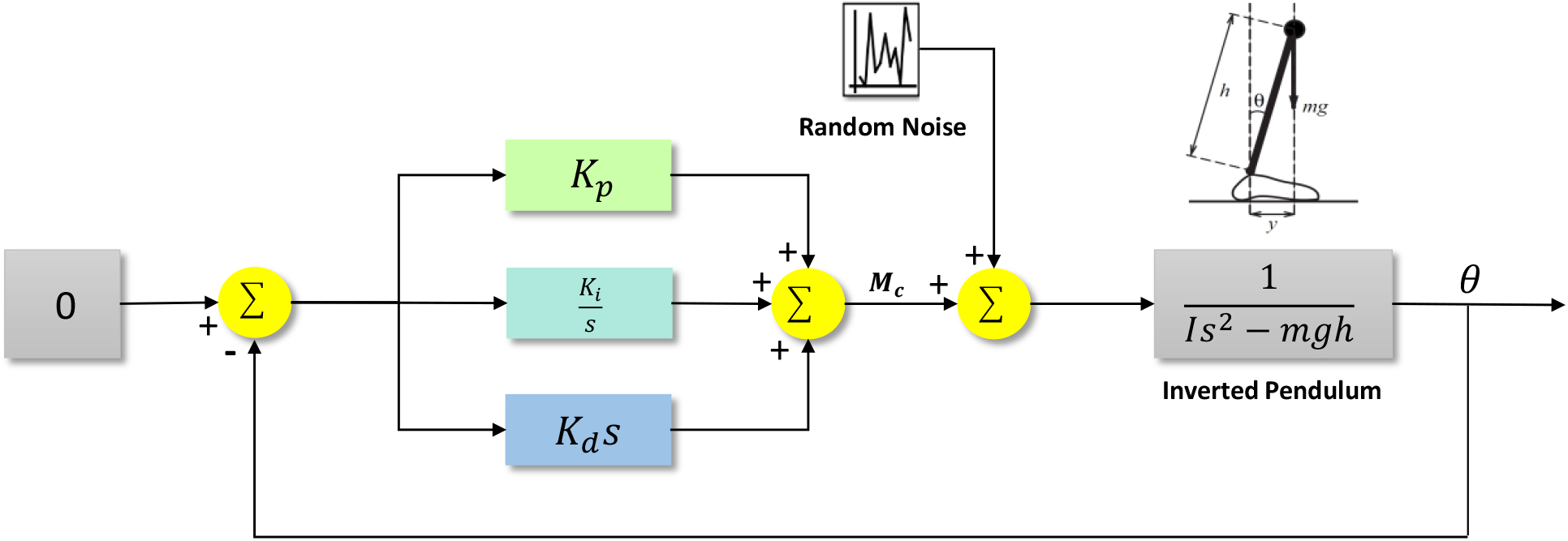
The simplified postural control block diagram and implemented in simulations

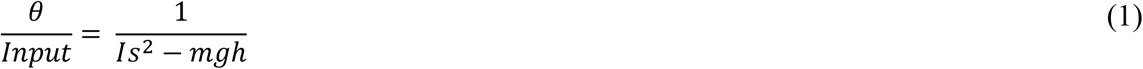

where *Input* = *M*_*c*_ +*Random noise,M*_*c*_ is a motor command that is an input to the body’s model, *Random noise* represents the summation of all internal noise inducing spontaneous body sway (Masani et al., 2006), *I* is the moment of inertia, *m* is the body weight, *g* is the gravitational acceleration, and *h* is the distance between CoM and the ankle. The output of the model is *θ*, the angle between the body and the vertical axis (body angle). Three blocks are for the PID controller (i.e., CNS) that regulates the plant (i.e., the body) (Eq. (2))

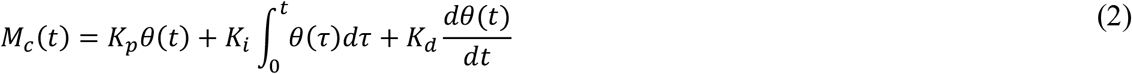

As mentioned before, *θ* is the body angle, 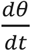 is the angular velocity of the body, and 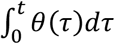 is the angular displacement of the body. *K*_*p*_ is usually corresponded to the joint impedance stiffness and *K*_*d*_ is represented the joint impedance viscosity in the model. We introduced the attention level as an integral coefficient (*K*_*i*_) in the PID controller, which demonstrates the role of CNS.

The value of *θ* is associated with posture balance control. That is, increasing the value of *θ* can negatively affect balance control. Therefore, the attention load should be introduced into the model in a way that by decreasing the focus of the attention to posture balance maintenance, the *θ* angle should be increased and vice versa. Attention should have a place in CNS. As mentioned before, CNS was modelled by the PID controller. The parameters of the PID controller (i.e., *K*_*p*_, *K*_*d*_, and *K*_*i*_) are associated with the gains that CNS implement to regulate balance. Since the proportional and differential coefficients of this controller have been reserved for stiffness and viscosity factors (Masani, Popovic, Nakazawa, Kouzaki, & Nozaki, 2003; Masani et al., 2006), we attributed *K*_*i*_ to the attention load. Therefore, we expected that increasing *K*_*i*_ can lead to the increment of the *θ* value (i.e., balance distribution).

The values of the models’ parameters were set based on the values reported in [16]. These values are reported in Table 1.

**Table 1.** The parameters of the model and their values

| Parameter | Description | Value |
| --- | --- | --- |
| m | Body weight | 76 Kg |
| I | Moment of inertia | 66 Kg m<sup>2</sup> |
| h | The distance of the COM to the ankle | 0.87 m |
| g | Acceleration of gravity | 9.8 m/s<sup>2</sup> |
| K<sub>d</sub> | Balance regulatory coefficient (viscosity) | * |
| K<sub>p</sub> | Balance regulatory coefficient (stiffness) | * |
| K<sub>i</sub> | Balance regulatory coefficient (attention) | * |
| * Different values were examined. |  |  |
| Variables | Description |  |
| $\theta$ | Body angle | |
| $\frac{d\theta}{dt}$ | Angular velocity | |
| $\int_0^t \theta(\tau) d\tau$ | Angular displacement | |
| $M_c$ | Motor command | |

## 3. Results

In the present study, the goal was to investigate the effect of attention in balance maintenance and introduce this effect into a postural balance control model. To do so, an experiment with two phases (single and dual-task) was performed. Both results of the experiment and a balance control model (considering the influence of the attention load) are discussed in this section.

### 3.1. Experiment results

The results of the experiment were studied considering the deviation of the wobble board (distance from the horizontal line) while performing the single task in comparison with the dual-task.

The mean of the wobble board distance from the horizontal line is reported in Figure 3. It can be seen that while carrying out the dual-task (posture balance maintenance while listening to audio numbers actively), the mean of wobble board deviations is more than that of the single task. This result is consistent with previous studies that referred to the limited capacity of the attention while performing a dual-task. In such cases, attention is dedicated to both tasks, and consequently, the performance in both tasks may be disrupted. In other words, the increment of the attention load in dual-task leads to the decrement of the balance control.

**Figure 3.**
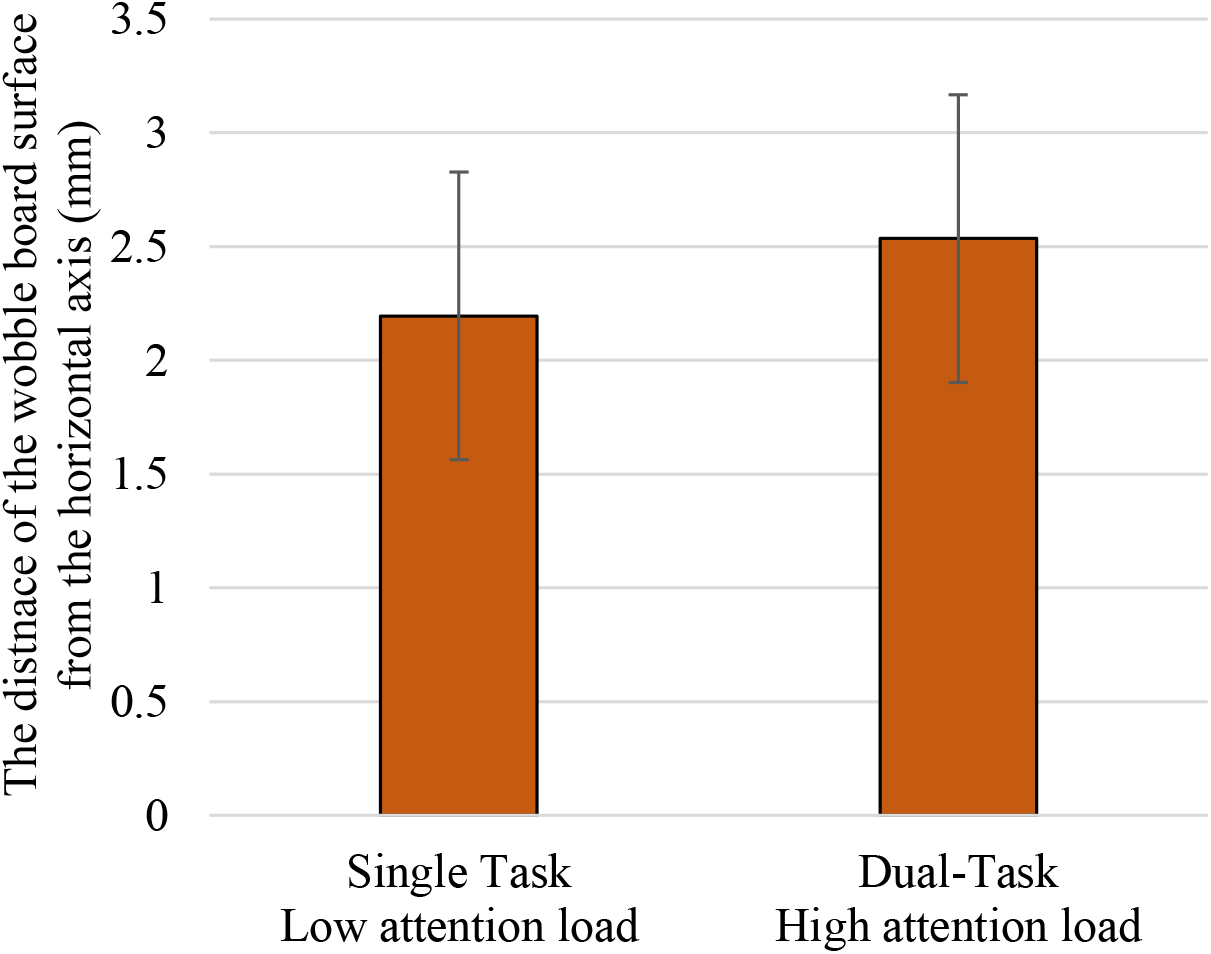
Comparison of the wobble board deviations between single and dual tasks; error bars indicate standard error of mean

### 3.2. Results of the model simulation

The proposed model was simulated in the MATLAB on the hardware of the LENOVO laptop with Intel® Core™ i7-4720HQ CPU @ 2.60 GH operation system. In simulations, we investigated the effect of *K*_*i*_ (attention load) on the angle between the body and the vertical axis, *θ* (posture imbalance) with different values of *K*_*p*_ and *K*_*d*_.

Figure 4 shows how increasing the value of *K*_*i*_ (i.e., the increment of the attention load) leads to the increment of the angle between the body and the vertical axis, *θ* (i.e., posture imbalance disruption).

**Figure 4.**
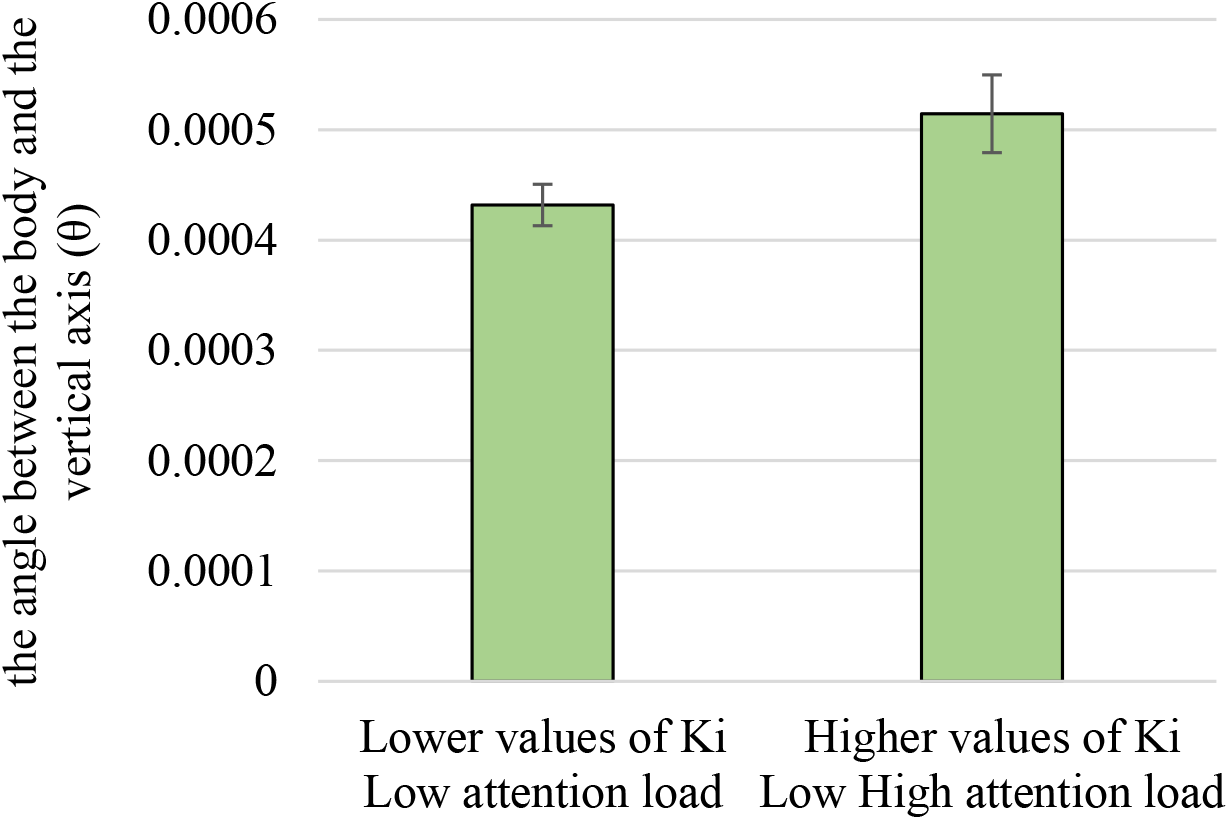
Comparison of the angle between the body and the vertical axis, *θ*, between two conditions 1-lower values of *K*_*i*_ (300-650), and 2-higher values of *K*_*i*_ (700-950), with different values of *K*_*p*_ and *K*_*d*_ (1000-1950)

Figure 5 shows the role of *K*_*p*_ and *K*_*d*_ in interaction with *K*_*i*_.

**Figure 5.**
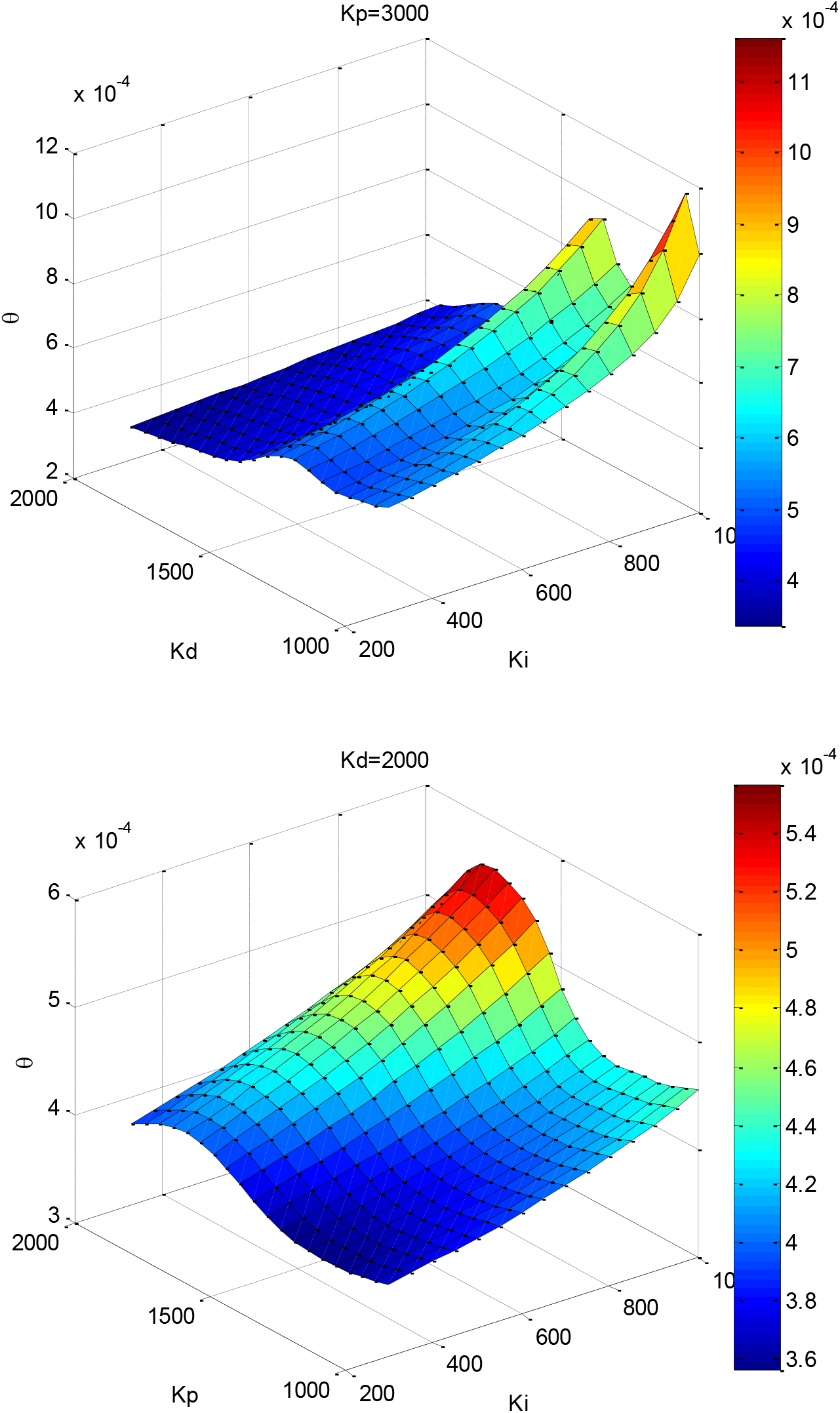
The values of the angle between the body and the vertical axis, *θ*, while the parameters *K*_*p*_, *K*_*d*_ and *K*_*i*_ are being varied; the top panel is for *K*_*p*_ = 3000 and the bottom panel is for *K*_*d*_ = *2000*.

As shown in Figure 5, increasing the value of *K*_*i*_ (the attention load) results in the increment of *θ* values. However, the level of increment is highly dependent on *K*_*p*_ and *K*_*d*_ values. The top panel of Figure 5 indicates that the slope of the increment of *θ* while increasing the *K*_*i*_ value in higher values of *K*_*d*_ (i.e., higher than 1500) is smaller than that of *K*_*d*_ in lower values (i.e., *K*_*d*_ is lower than 1500). Conversely, the bottom panel of Figure 5 demonstrates that the slope of the increment of *θ* while increasing the *K*_*i*_ value in lower values of *K*_*p*_ (i.e., lower than 1500) is smaller than that of *K*_*p*_ in higher values (i.e., *K*_*d*_ is higher than 1500). Simulations also revealed that for some particular values of *K*_*p*_ and *K*_*d*_ (approximately lower than 250), regardless of *K*_*i*_ value, the system is not stable.

## Discussion

The impact of attention disruption on postural balance control was investigated experimentally in the present study. This effect was also modelled computationally by introducing the attention load to a previously announced balance control model.

The results of the experiment show that increasing the attention load in a dual-task in comparison with performing a single task disturbs the balance control. This result is in line with the claim of attention capacity limitation in individuals (Just & Carpenter, 1992; McDowd, 2007; Woollacott & Shumway-Cook, 2002). According to this limitation, at times when various processing systems (here, the system for response to auditory stimuli and balance controller) compete for neuronal resources, the amount of attention allocated to each processing system reduces. Reduction of attention results in either balance control disruption or cognitive performance impairment. Therefore, it can be claimed that besides joint impedance stiffness and viscosity, the attention load is critical for balance control maintenance. These influential factors were introduced to the proposed models as the factors of a PID controller. Comparing the results shown in Figure 3 and Figure 4 demonstrates that the behaviour of the proposed model is in agreement with the experiment outcomes.

Scrutinizing the results shown in the top panel of Figure 5 reveals that in the lower values of *K*_*p*_ (i.e., less than 1500), changing the attention load has no considerable effect on the posture balance control (i.e., the value of *θ*). However, by increasing the stiffness factor (*K*_*p*_), the role of attention load increases. In other words, when more weight is put on the stiffness factor, less attention load (more attention level) is required to keep the *θ* value small (i.e., balance control maintenance). From a control engineering perspective, increasing the value of *K*_*d*_ puts more weights on the derivative element in the PID controller. Putting more weights on the derivative control increases the sensitivity of the system to noise and disturbances (Johnson & Moradi, 2005; Saab, 2017; Sekara & Matausek, 2009). Therefore, in our study, it is expected that less attention load should be implemented to overcome the disturbance sensitivity. This expectation is in agreement with the results shown in the top panel of Figure 5. In studies on motor control, it was also shown that lower levels of joint impedance viscosity could lead to higher oscillations in the joint angle (Farzad Towhidkhah, 1998; F Towhidkhah, Gander, & Wood, 1997). In those previous studies, the role of attention has not been considered. The outcomes of the current study demonstrate that the claim of increasing joint angle oscillations due to the lower level of joint impedance viscosity is not always valid and depends on the attention load. However, in higher values of joint impedance viscosity, the sensitivity of the system to the attention load decreases (see the top panel of Figure 5 for *K*_*d*_ *>* 1500).

The bottom panel of Figure 5 shows that in lower values of joint impedance stiffness (*K*_*p*_ *<* 1500), the sensitivity to the attention load is lower than the high values. Therefore, in high values of joint impedance stiffness and low values of joint impedance viscosity, the sensitivity of the balance control system (the *θ* value) to the attention load is higher than other conditions. Figure 5 also demonstrates that the issues created for the joint angle -due to the improper amount of joint impedance stiffness -can be compensated by changing the attention load. Since fatigue, exercises, or other motor-control related activities affect the level of viscosity and stiffness (Chalchat et al., 2020; Racinais, Cocking, & Périard, 2017), how these two factors interact with the attention factor should be considered more carefully in sports and rehabilitation activities.

The model’s simulation results reveal that there is a bell-shaped relationship between stiffness (*K*_*p*_) and balance control. That is, increasing stiffness property leads to balance disruption (i.e., increasing the *θ* value), at first, but more increment gradually leads to better balance control (i.e., lower values of *θ)*. Such a bell-shaped relationship is typical in biological phenomena (Cardinal, 2015; Fernandez-de-Cossio-Diaz & Vazquez, 2018; Radak et al., 2017). This result suggests that the claim of the enhancement of balance control by the increment of stiffness is not always valid and needs further inspection. These outcomes can be useful for researchers and users in sports that need to maintain balance. In that, a coach or a researcher in the field of sports should pay attention to the fact that the level of focus on maintaining balance should be set to an optimal value considering both joint impedance stiffness and viscosity.

## Acknowledgment

The authors would like to acknowledge the Amirkabir University of Technology for its support and providing the data acquisition facility.

## Funding Sources

This research did not receive any specific grant from funding agencies in the public, commercial, or not-for-profit sectors.

## Availability of data and material

The recorded data (except for the participants’ personal information) can be available by sending the request to the first or the corresponding author.

## Declaration of interests

None of the authors has a financial interest or conflict of interest.

## Appendix

**Table 1.** A brief review of several studies that investigated the effect of attention on balance control

| No. | Balance criteria | Cognitive criteria | Balance task | Cognitive task | Age | Subjects | Result | Ref. |
| --- | --- | --- | --- | --- | --- | --- | --- | --- |
| 1 | COP moving window | Look straight | Stance | Focus on lessening the leg movements | Mean age of 56 and 61 | People who have done hemotherapy + the elderly | No difference in balance control | (Arnold, 2014) |
| 2 | The time the platform has stayed horizontal | Look straight | Stance | Focusing on keeping the platform horizontal | 86-60<br>With a mean age of 69.4 | Healthy elderly | External attention helps to keep balance better | (Chiviacowsky, Wulf, & Wally, 2010) |
| 3 | CoP characteristics | Right answers | Stance and balance shift to protheses | Announce whether the equations are right or wrong | 23-78 mean: 59.4<br>Control: Mean 58.9 | Amputee people | More wrong answers of amputees + poor balance of amputees + better learning of amputees | (Geurts & Mulder, 1994) |
| 4 | Computerized Dynamic Posturography | Equilibrium score + Sensory ration + Strategy Score | Stance | Sensory organization test | children | ADHD + control | Poor balance in ADHD children | (Ren et al., 2014) |
| 5 | CoP and CoM characteristics | Reaction time + announcement accuracy | Sit (dual-task) + walk (dual-task) | Announce the tone of the sound (High/low) | Patients: mean: 21<br>Controls: 20.7 | People with concussion + controls | Lower COM velocity in concussed people | (Catena, van Donkelaar, & Chou, 2011) |
| 6 | CoP characteristics | Number of right answers | Stance | Memorize the played songs | Children | ADHD + control | More fluctuations in ADHD children | (Shorer et al., 2012) |
| 7 | CoP position | The right number of shopping list items | 5 mins walking + stance | Memorize the shopping list items | Patients: Mean: 66<br>Control: Mean: 62.3 | People with stroke + control | A better balance of the control group | (Hyndman et al., 2006) |
| 8 | Balance master, Computerized posturography | The time spent on balance correction | Stance | Brook test | 18-23 | Females | More difficult balance tasks affected cognitive task | (Swan, Otani, & Loubert, 2007) |
| 9 | Balance score | Symbol digit modalities test | Berg balance scale | SDMT | Mean: 39.4 | Acquired brain injury people | In dual-task lowering the speed and accuracy were observed. | (McCulloch, Buxton, Hackney, & Lowers, 2010a) |
| 10 | Ankle and gate characteristics | Speed and accuracy of the answers, oral time answering | Walking | Reverse counting, stating items' names, auditory Stroop test | More than 65 | The elderly with balance control difficulties | Lowering the angle in $VP^{50}$ in comparison with $FT^{51}$ and $ST^{52}$ | (Silsupadol et al., 2009) |

## Notes

### Competing Interest Statement

The authors have declared no competing interest.

